# BiomeSeq: A Tool for the Characterization of Animal Microbiomes from Metagenomic Data

**DOI:** 10.1101/800995

**Authors:** Kelly A. Mulholland, Calvin L. Keeler

**Author notes:** Corresponding author: Department of Animal and Food Sciences, University of Delaware, Newark, DE 19716.

## Abstract

The complete characterization of a microbiome is critical in elucidating the complex ecology of the microbial composition within healthy and diseased animals. Many microbiome studies characterize only the bacterial component, for which there are several well-developed sequencing methods, bioinformatics tools and databases available. The lack of comprehensive bioinformatics workflows and databases have limited efforts to characterize the other components existing in a microbiome. BiomeSeq is a tool for the analysis of the complete animal microbiome using metagenomic sequencing data. With its comprehensive workflow, customizable parameters and microbial databases, BiomeSeq can rapidly quantify the viral, fungal, bacteriophage and bacterial components of a sample and produce informative tables for analysis. BiomeSeq was employed in detecting and quantifying the respiratory microbiome of a commercial poultry broiler flock throughout its grow-out cycle from hatching to processing. It successfully processed 780 million reads, of which 5,163 aligned to avian DNA viral genomes, 71,936 aligned to avian RNA viral genomes, 469,937 aligned to bacterial genomes, 504,682 aligned to bacteriophage genomes and 1,964 aligned to fungal genomes. For each microbial species detected, BiomeSeq calculated the normalized abundance, percent relative abundance, and coverage as well as the diversity for each sample. BiomeSeq provides for the detection and quantification of the microbiome from next-generation metagenomic sequencing data. This tool is implemented into a user-friendly container that requires one command and generates a table consisting of taxonomical information for each microbe detected as well as normalized abundance, percent relative abundance, coverage and diversity calculations.

## Background

Specific and unique animal microbiomes contribute to the biological function of various locations on the body including the gut, skin, vagina, oral cavity, and respiratory tract (Cui *et al.*, 2013). Disturbances of these environments by colonization of a new bacteria, eukaryotic virus, or fungi can lead to competition, invasion and replacement. Under appropriate conditions this may result in disease. Advancements in next-generation sequencing technology enable investigations into individual components of the microbiome, thereby gaining insight into the dynamic interactions taking place (Barzon *et al.*, 2011). Identification of microbial communities within these environments can aid in elucidating the role they play in both healthy and diseased animals.

Recent studies attempting to characterize the microbiomes of mammals have focused primarily on their bacterial composition, as there are well established and rapid methods of sequencing and analyzing this component (Bond *et al.*, 2017; De Boeck *et al.*, 2015; Gaeta *et al.*, 2017; Glendinning *et al.*, 2017; Johnson *et al.*, 2018; Shabbir *et al.*, 2015). The 16S rRNA gene is commonly used to identify and compare the bacterial genera present in a given sample (Clarridge *et al.*, 2004). Accessible bacterial databases, such as Greengenes (De Santis *et al.*, 2006) and Silva (Quast *et al.*, 2013), in addition to well-developed bioinformatics workflows are available to facilitate these analyses (Meyer *et al.*, 2008; Caporaso *et al.*, 2010; Schloss *et al.*, 2009). Internal Transcribed Spacer, or ITS, is a widely used fungal genetic marker gene. Similar to 16S rRNA, accessible fungi databases (Kõljalg *et al.*, 2013) and bioinformatics workflows for fungal analysis exist (Caporaso *et al.*, 2010).

Characterizing the viral component of the microbiome presents unique difficulties. Unlike the ribosomal genes of bacteria and fungi, viruses are heterogeneous in their genetic content and therefore do not have a conserved genomic region that can be easily sequenced and employed for taxonomic classification (Zou *et al*., 2016). In addition, host DNA contamination has been found to negatively impact the interpretation of results (Daly *et al.*, 2015). As a result, there have been fewer efforts to develop comprehensive viral genome databases similar to those available for bacteria (De Santis *et al.*, 2006; Quast *et al.*, 2013; Kõljalg *et al.*, 2013). Quantification of viral abundance is another limitation with characterizing the virome. Due to the lack of eukaryotic viral genome databases, a sequence-similarity independent approach is often employed to detect eukaryotic viruses, but this approach does not allow for accurate abundance calculations. In addition, many of the available virome bioinformatics tools require the user to possess extensive command-line knowledge and computational resources to successfully install and run the necessary programs and their dependencies on the command line. A user-friendly tool for the analysis of the viral, fungal, bacterial and bacteriophage components is essential to elucidating the complete ecology of a microbiome.

Herein, we present BiomeSeq, a tool for the analysis of complete animal microbiomes from metagenomic data. The BiomeSeq workflow and databases address the challenges of characterizing the eukaryotic virome by including quality filtering and host decontamination, sequence-similarity dependent alignment to microbial reference genome databases and accurate quantification of microbial abundance. It also analyzes the fungal, bacteriophage and bacterial components using the same sequencing data to produce a complete analysis of the microbiome without requiring additional sequencing of the 16S rRNA and ITS genes. Additionally, utilizing shotgun metagenomics to analyze the bacterial and fungal components can increase taxonomic resolution, permit analysis of complete genomes instead of a conserved genomic region, and allow for a comparison of bacteria and fungi to the viral and bacteriophage components (Jovel *et al.*, 2016). BiomeSeq is available as a user-friendly docker container. This versatility allows BiomeSeq to be accessible to users with varied degrees of command-line knowledge and computational resources. While BiomeSeq has been developed and tested on avian species, it can be used to characterize microbiomes of a variety of species.

## Implementation

BiomeSeq is currently available as an open-access and user-friendly tool on Docker Hub. As the docker container is self-contained, it simplifies installation and execution by eliminating the need for downloading and installing dependent software and requires only one command. Additionally, BiomeSeq is customizable and allows the user to adjust parameters similar to a command-line tool. Table 1 includes all software and parameters used in BiomeSeq.

**Table 1.** Software tool and parameters used in BiomeSeq

| **Process** | **Tool Name** | **Parameters** |
| --- | --- | --- |
| Quality | Trim Galore |  |
| Decontamination | Bowtie 2 | -x -S |
|  | Samtools | view -bS |
| Microbial Database Alignment | Bowtie 2 | -x -S |
|  | Samtools | view -bSq [user input] |

BiomeSeq accepts both single- and paired-end reads in fastq format generated by DNA-seq or RNA-seq methods. Along with the fastq file, the user may customize a number of parameters including: the host genome that the sample was derived from, a host-specific viral database, mapping quality threshold, output file name and an output directory. Figure 1 shows an overview of the BiomeSeq workflow. BiomeSeq generates a table consisting of NCBI RefSeq accession number, microbe name, taxonomy, number of mapped reads in the file, normalized abundance, percent relative abundance, genome coverage for each eukaryotic virus, bacteria, bacteriophage and fungi detected, as well as an alpha diversity calculation for the sample. Table 2 is an example of an output table for the viral component. Similar tables are generated for bacteria, bacteriophage and fungal data. Visualizations of this data can be easily generated using several different packages in R.

**Table 2.** Example table generated by BiomeSeq of the viral component of a commercial poultry flock at Week 6.

| Ref Seq Number | Name | Taxonomy | Genome Size | Number Mapped | Norm. Abundance | Relative Abundance | Genome Coverage | Diversity |
| --- | --- | --- | --- | --- | --- | --- | --- | --- |
| NC002229 | Gallid Alphaherpesvirus 2 | Double Stranded; Enveloped; Herpesviridae; Mardivirus | 177874 | 1 | 27.38 | 0.10% | 0 | 0.534 |
| NC002577 | Gallid Alphaherpesvirus 3 | Double Stranded; Enveloped; Herpesviridae; Mardivirus | 164270 | 1 | 29.65 | 0.10% | 0 |  |
| NC015396 | Avian Gyrovirus | Single Stranded; Non-Enveloped; Circoviridae; Gyrovirus | 2383 | 72 | 147165.92 | 15.47% | 3.05 |  |
| NC001720 | Fowl Aviadenovirus | Double Stranded; Non-Enveloped; Adenoviridae; Aviadenovirus | 43804 | 4560 | 507048.92 | 53.30% | 10.51 |  |

**Figure 1.**
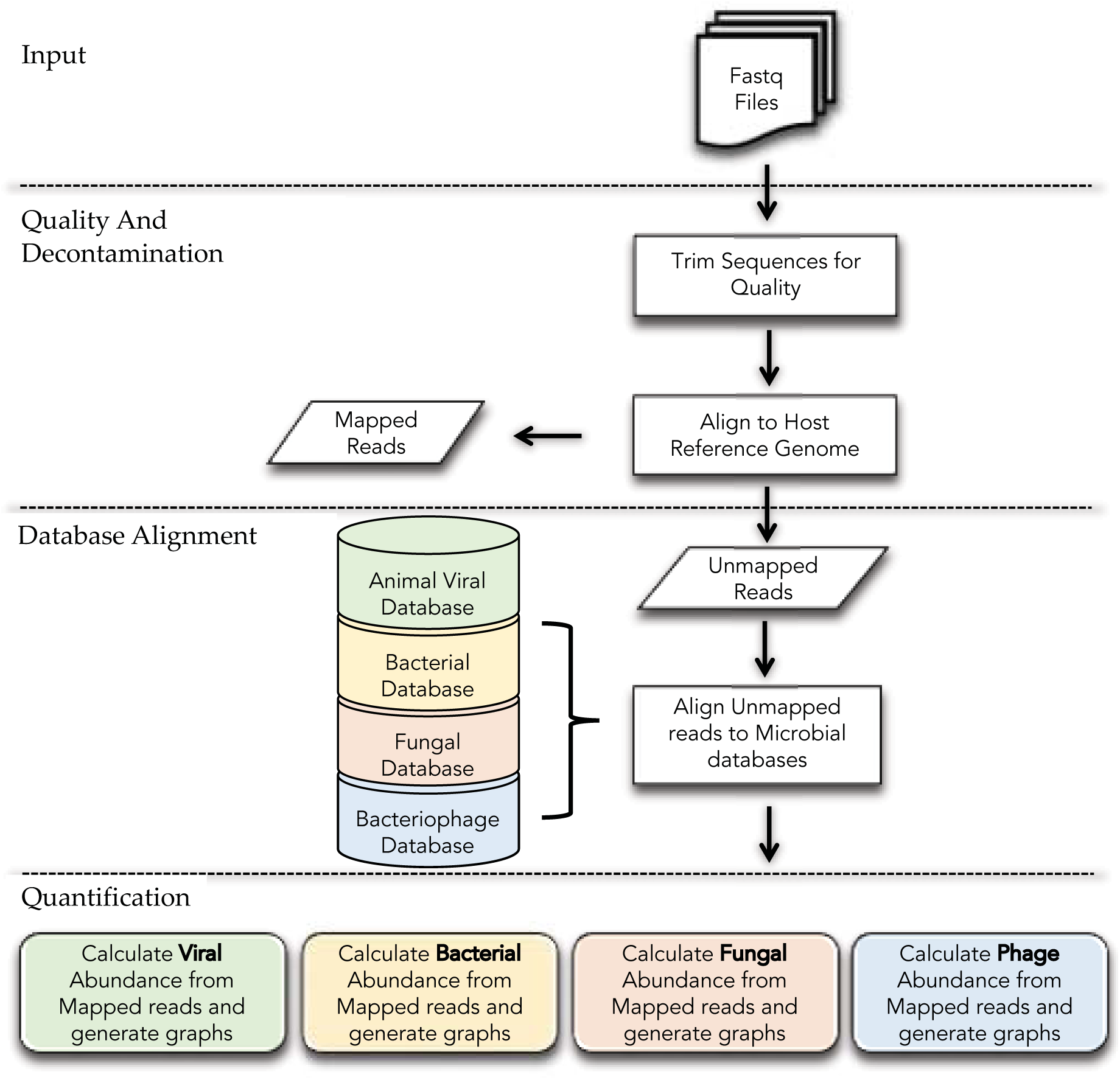
BiomeSeq Workflow.

### Quality and Decontamination

Individual fastq sequence files are first analyzed for per-base sequence quality, per-sequence quality, sequence length distribution, duplicate sequences, and overrepresented Kmers. Reads with a quality phred score below 30, reads under 100 base pairs and adapter sequences are removed. Quality steps are processed using Trim-Galore (Martin *et al.*, 2011). The remaining reads are then aligned to the user-specified host reference genome using Bowtie2 and only unmapped reads are extracted and analyzed further (Langmead *et al.*, 2012). This step removes host genome contamination from the data, increasing analytical efficiency and mapping accuracy (Daly *et al*., 2015).

### Databases

The remaining sequencing reads are aligned to an host-specific viral genome database, a bacterial database, a fungal database and a bacteriophage database using the Bowtie 2 alignment algorithm (Langmead *et al.*, 2012). Mapping quality threshold default is 20, however this parameter may be customized by the user. The eukaryotic viral genome database currently includes avian-specific viral genomes and was constructed using full genome reference sequences of both DNA and RNA avian viruses obtained from the National Center for Biotechnology Information (NCBI) Virus Database (O’Leary, 2016). The avian DNA viral genomes include 48 viral elements from 9 unique families and the avian RNA viral genomes include 63 viral elements from 13 families. The avian DNA and RNA viral database is organized by the classification of their viral structure and genome organization. DNA viruses are organized hierarchically by whether the virus is double- or single-stranded and whether the virus is enveloped or non-enveloped. RNA viruses are organized hierarchically by whether the virus is double- or single-stranded, negative or positive sense, segmented or non-segmented and whether the virus is enveloped or non-enveloped. The eukaryotic viral genome database will include additional host-specific viruses from a variety of species.

Custom bacterial, fungal and bacteriophage databases were constructed using complete and representative genomes obtained from the NCBI Reference Sequence Database and contain 3,623, 1,281 and 2,212 genomes, respectively (O’Leary, 2016). Each microbial database and corresponding aligner index files can be downloaded from CyVerse. As an additional feature, BiomeSeq also accepts custom microbial databases provided by the user.

### Quantification and Output

A sequence similarity-dependent approach for detecting viruses contributes to the rapid detection of known viruses while also allowing for the quantification of biodiversity, which similarity-independent approaches lack (Herath *et al*., 2017). This approach can be applied to bacteria, fungi and bacteriophage as well. For each individual sample, the reads that map to each microbe are normalized based on both microbial and reference genome length per 100,000 host cells using an adaptation to the equation presented by Moustafa and his colleagues in 2017 to quantify viral abundance (Moustafa *et al.*, 2017):

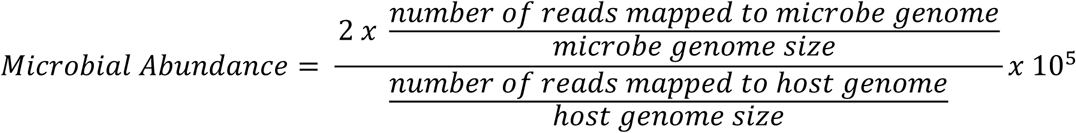

Percent relative abundance is quantified using the following equation:

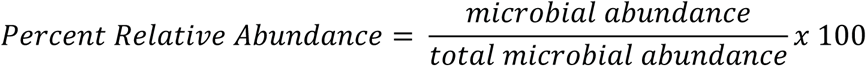

Genome coverage is approximated using the following equation:

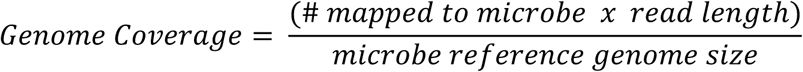

Alpha diversity for each sample is calculated using the Shannon Diversity Index, a commonly used equation for calculating species diversity in a microbiome as it accounts for both abundance and evenness of the species.

## Results and Discussion

### A Longitudinal Study of the Microbial Ecology of a Healthy Broiler Flock

Recent studies have identified specific bacterial and viral agents within the respiratory microbiome of both humans and animals that are associated with the severity and spread of disease (Bakaletz *et al.*, 1995; Pettigrew *et al.*, 2008; De Steenhuijsen Piters *et al.*, 2016; Teo *et al.*, 2015). In order to understand the complex etiology of a disease and the changes in the microbial ecology of a diseased microbiome, a comprehensive analysis of a healthy microbiome is first required. BiomeSeq was employed to detect and quantify eukaryotic viruses, bacteria, bacteriophage, and fungi in a healthy commercial broiler flock during the grow-out cycle from hatching to processing.

Tracheal swabs were collected at hatching and at weekly intervals through processing at day 50 (8 samples) from an antibiotic-free commercial broiler flock. Both DNA and RNA were isolated and sequencing was performed for each of the eight time points using the Illumina HiSeq platform producing 1 × 100 single-end reads. Each of the resulting 16 samples were processed using BiomeSeq with the following parameters: -g chicken -d avianALL_db -q 20.

In total, BiomeSeq detected 5,163 reads aligned to avian DNA viruses and 71,936 reads aligned to avian RNA viruses. A total of 11 viral species, representing 9 genera and 8 families, were identified from the avian respiratory tract during the grow-out period. This data is represented in a heatmap (Figure 2). A total of 469,937 reads were aligned to the bacterial genome database. A total of 533 unique bacterial species were identified, of which 45 had a calculated relative abundance greater than 0.5%. The 45 most abundant species detected extend from 4 phyla, 7 classes, 13 orders, 26 families and 45 genera. This data is represented in a phylogenetic tree generated using the Phytools package in R (Figure 3; Revell, 2012). A total of 504,682 reads aligned to the bacteriophage genome database. A total of 30 unique bacteriophage species extended from 1 classified and 1 unclassified order, 4 classified and 1 unclassified families, and 5 classified and 4 unclassified genera were identified. This data is represented in a Venn diagram of the common bacteriophage species detected at Week 0, Week 3 and Week 7, generated using the VennDiagram package in R (Figure 4; Chen *et al.*, 2011). A total of 1,964 reads aligned to the fungal genome database. Sixty-one unique fungal species were identified which extended from 2 phyla, 9 classes, 20 orders, 37 families and 50 genera. This data is represented in a microbial network generated with Cytoscape (Figure 5; Shannon *et al*., 2003).

**Figure 2.**
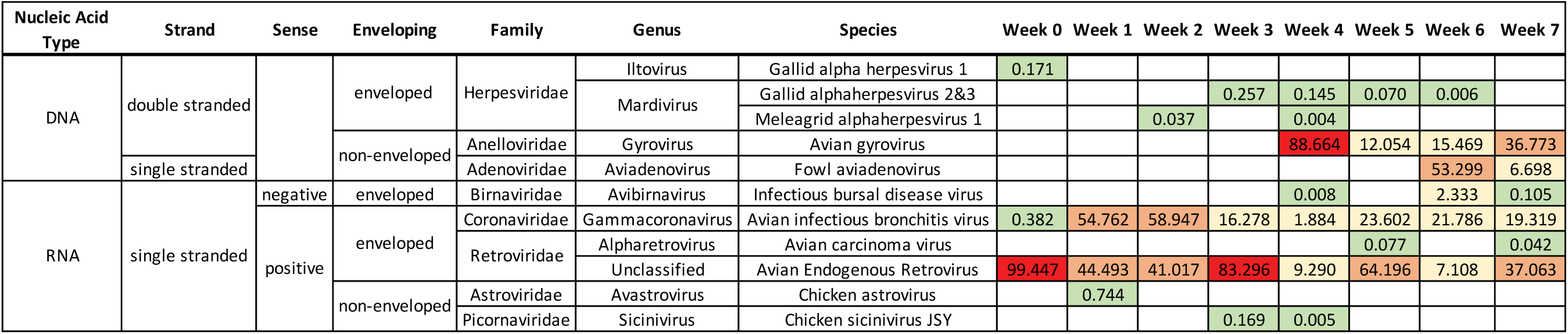
Heatmap consisting of each virus species identification and abundance in a commercial poultry flock from hatching to processing. Color corresponds to the range of relative abundance of each family from 0 to 100%. Green: 0-1%; yellow: 1-25%; orange: 25-75%; and red: 75-100%. The sum of each column, or week, is 100%.

**Figure 3.**
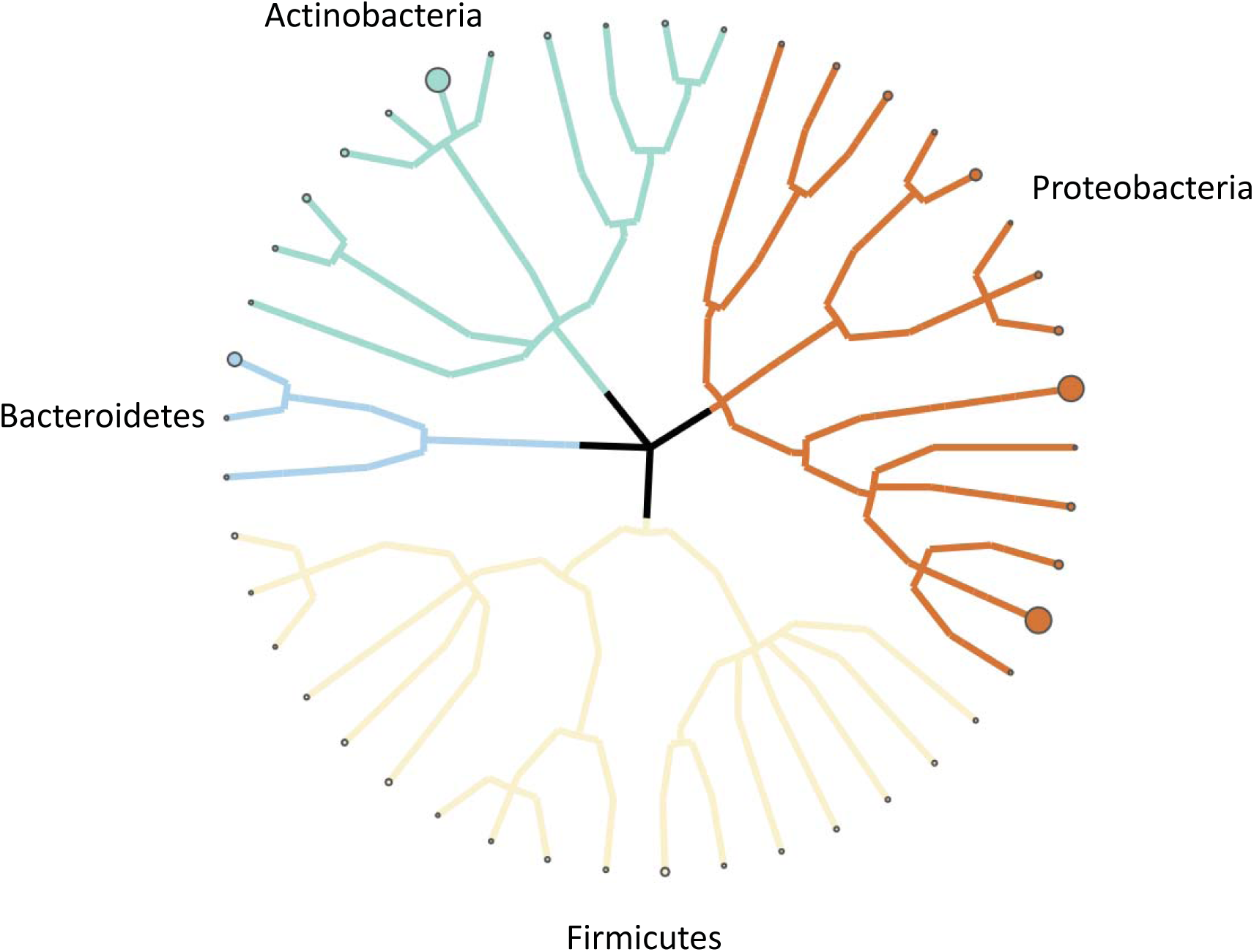
Phylogenetic tree of bacterial species detected in a commercial poultry flock. Branches extend from phylum to species. Nodes indicate detected species and diameter indicates average abundance.

**Figure 4.**
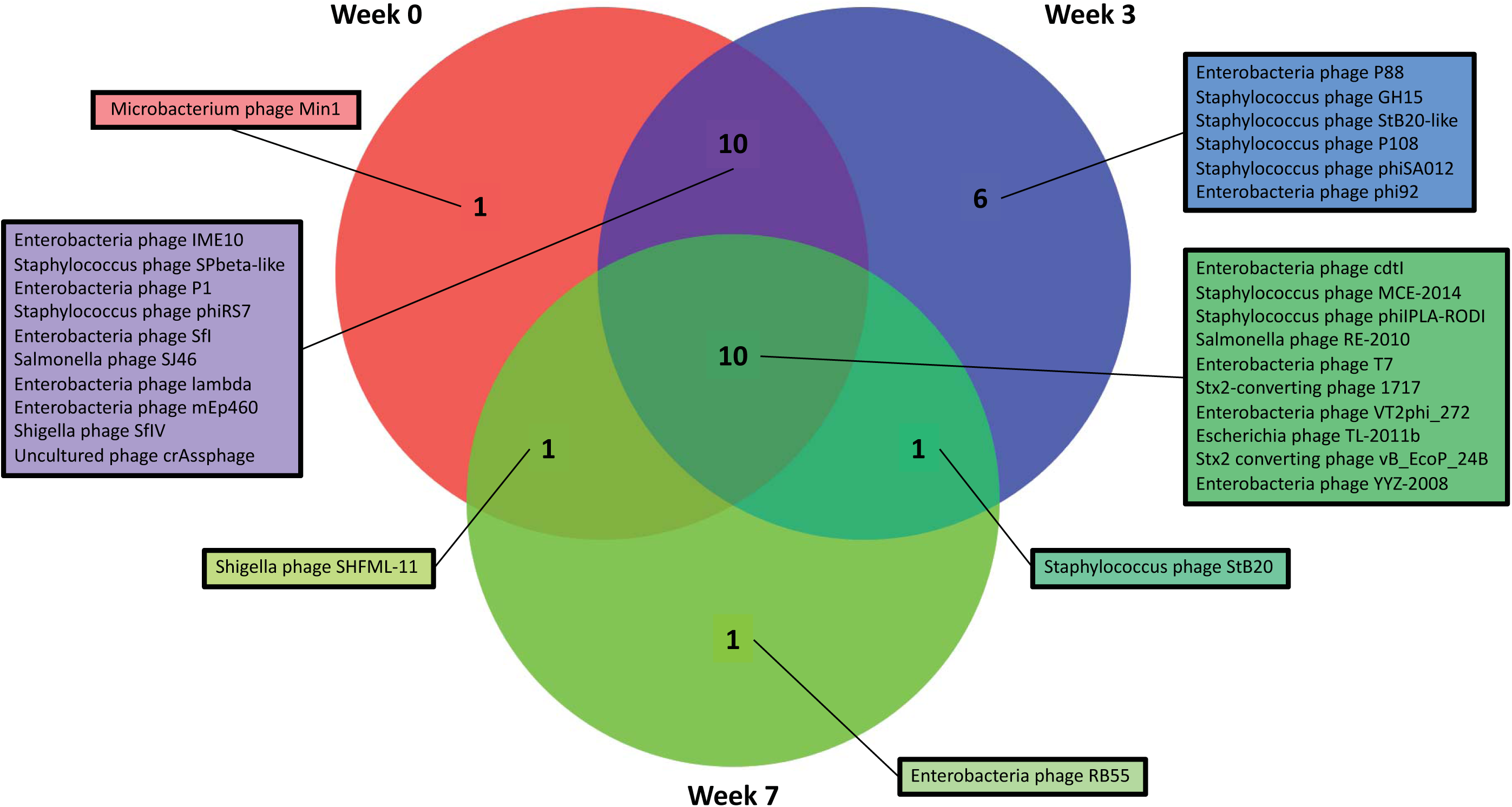
Venn Diagram of the detected bacteriophage species in a commercial poultry flock at Week 0, Week 1 and Week 7.

**Figure 5.**
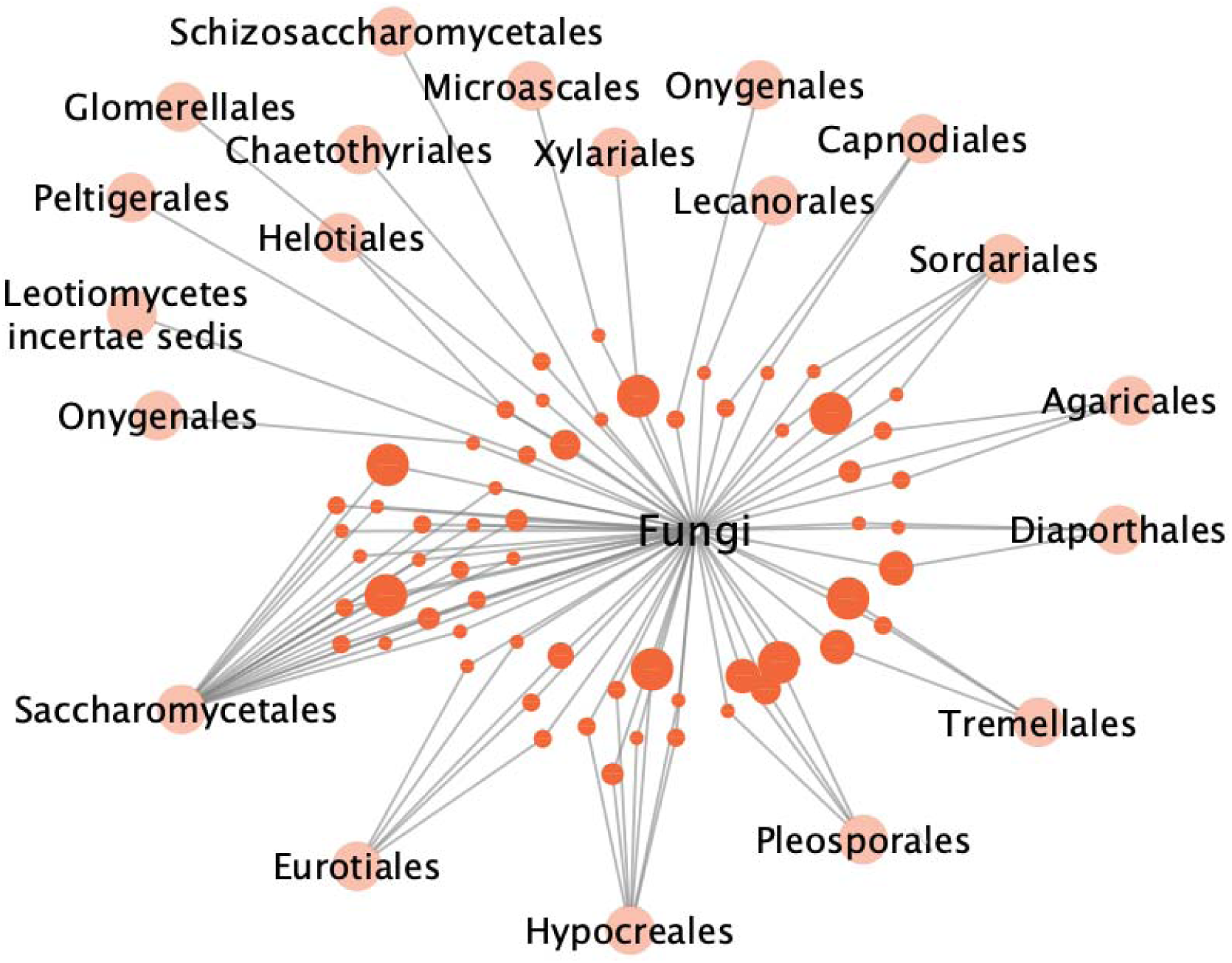
Network of fungal species detected in a commercial poultry flock. Outer nodes represent order level, while inner nodes represent species. Diameter of the inner nodes correlate to species frequency, or the number of weeks the species was detected.

## Conclusions

The complete characterization of a microbiome is critical in elucidating the complex ecology of the microbial composition within healthy and diseased animals. Recent studies have focused on the bacterial component, as there are several well-developed sequencing methods, bioinformatics tools and databases available. The lack of comprehensive bioinformatics workflows and databases have limited efforts to characterize the other components. BiomeSeq is a tool for the analysis of the animal microbiome using metagenomic data. With its comprehensive workflow and custom databases, this tool can rapidly quantify the eukaryotic viral, fungal, bacteriophage and bacterial components of a sample and produces informative tables for analysis. The sequence-dependent approach that BiomeSeq utilizes provides the necessary information required to accurately quantify microbial abundance, genome coverage and diversity. Conversely, this method limits BiomeSeq’s ability to perform in de novo microbe discovery. Moreover, a sequence-dependent approach including only representative microbial genomes may underrepresent the abundance of specific microbial strains. To resolve these limitations, BiomeSeq accepts custom microbial databases provided by users which may include microbial genomes derived by other host species and novel microbial sequences.

BiomeSeq was employed in detecting and quantifying the respiratory microbiome of a commercial poultry broiler flock throughout its grow-out cycle from hatching to processing. This study provides the first comprehensive analysis of the ecology of the avian respiratory microbiome and will facilitate future investigations of animal diseases. BiomeSeq is accessible as a container, available as a user-friendly container on Docker Hub.

## Abbreviations

NGS: Next generation sequencing
16S rRNA: 16s ribosomal RNA
NCBI: National Center for Biotechnology Information
DNA: Deoxyribonucleic acid
RNA: Ribonucleic acid

## Declarations

### Ethics Statement

Not applicable.

### Consent for publication

All authors have consented to publication

### Availability of data and material

The BiomeSeq Docker container is available at http://dockerhub.com.

BiomeSeq custom databases are available at https://de.cyverse.org.

### Authors’ contributions

KAM is the primary author of this manuscript. KAM developed the bioinformatics workflow and constructed the animal-specific viral genome database, the fungal database, the bacteriophage database and the bacterial database. KAM wrote all programs for microbial calculations and programs to generate visual representations of microbial data. CLK is the corresponding author of this work. CLK contributed to the design of the work, the acquisition of samples, the analysis and interpretation of the data, and revised and edited the manuscript.

## Acknowledgements

We thank Monique Robinson, Sharon Keeler, Hong Li and Daniel Bautista for their contributions in collecting and processing experimental samples and Shawn Polson for his insightful comments and suggestions for this manuscript.

